# MYB93 regulates responses to environmental sulphur in *Arabidopsis* and tomato

**DOI:** 10.1101/2025.03.04.641362

**Authors:** Xulyu Cao, Helen B. Wilkinson, Bethany Hutton, Nancy McMulkin, Neil S. Graham, Ross Etherington, Alice Oliver, Clare M. Clayton, Harjeet Kaur, Juliet C. Coates

**Affiliations:** College of Resources and Environment, Academy of Agricultural Sciences, Key Laboratory of Efficient Utilization of Soil and Fertilizer resources, Southwest University, Chongqing, 400716, China; School of Biosciences, University of Birmingham, Birmingham B15 2TT, UK; Plant and Crop Sciences Division, School of Biosciences, Sutton Bonington Campus, University of Nottingham, LE12 5RD, UK

**Keywords:** MYB transcription factor, Sulphur, Root development, transcriptome

## Abstract

Sulphur (S) is an important nutrient that has wide-ranging effects on plant health and metabolism. Several classes of transcription factor respond to S deprivation, including R2R3-MYBs. In *Arabidopsis*, the *AtMYB93* transcription factor-encoding gene is upregulated by S deprivation. *At*MYB93 has a non-redundant function in lateral root development and redundant functions in suberin biosynthesis alongside related MYB transcription factors, but *At*MYB93’s role in S signalling, and how it relates to lateral root development, is unknown.

We show that the transcriptome of *Atmyb93* mutant roots implicates *At*MYB93 in responses to S, including changes in S transport and metabolism, and flavonoid- and carbohydrate metabolism. Elemental analysis demonstrates that the *Atmyb93* mutant has elevated shoot S levels while tomato *Sl*MYB93-overexpressing plants have reduced shoot S. We uncover a stimulatory effect of S deprivation on adventitious root development. However, *Atmyb93* mutants do not show significant changes in sensitivity to S with respect to lateral-or adventitious root development, most likely due to some functional redundancy. Moreover, *AtMYB93* promoter activity is not spatially regulated by S deprivation. Taken together, our data suggest that *AtMYB93* has a role in mediating root responses to S in alongside other root transcription factors.

## Introduction

Plants require a range of nutrients to grow and develop optimally. The soil provides macronutrients and micronutrients, in relatively large and small amounts respectively. The key macronutrients that plants obtain from soil are nitrogen (N), phosphorus (P), potassium (K), calcium (Ca), magnesium (Mg) and sulphur (S). S is often considered a ‘neglected’ macronutrient: S fertilisation can improve plant yield, abiotic- and biotic stress tolerance (Ali et al. 2021; Narayan et al. 2023; Sarda et al. 2013; Zenda et al. 2021) although S deprivation may improve resistance to some pathogens (Criollo-Arteaga et al. 2021). In terms of human nutrition, plants are the major source of methionine, an essential S-containing amino acid (Kopriva et al. 2019).

Problems with S deficiency in crops are increasing globally due to the decreasing amount of rainfall-derived S in combination with the continued development of high-yielding crop varieties (Sharma et al. 2024) (Zenda et al. 2021) (Sarda et al. 2013) (Grant et al. 2012). Crop requirements for S vary, with Brassicas (especially oilseed rape) requiring high S, cotton and sugarcane requiring medium S and cereals having relatively low S requirements (Zenda et al. 2021). However, naturally high S *Brassica* crops used for animal feed can promote risk of goitre and kale anemia (Smith 1980; Smith et al. 1974; Paxman and Hill 1974). Moreover, S levels can affect crop quality, for example high S can cause increased acrylamide formation in cooked potatoes (Elmore et al. 2007; Elmore et al. 2010). In marine ecosystems, sulfide stress in seagrass beds, caused by microbial anaerobic digestion of sulfate in sediments, increases with elevated temperature (Zhang et al. 2024). Thus, S levels in plants may require careful management on a case-by-case basis.

S deficiency in plants has wide-ranging effects including reduced chlorophyll, changes in sugar- and amino acid metabolism, changes in phenylpropanoid metabolism (particularly flavonoids) and changes in antioxidant levels (e.g. (Lunde et al. 2008; Wawrzynska et al. 2022; Bielecka et al. 2014; Chandra and Pandey 2014; Forieri et al. 2017; Henriquez-Valencia et al. 2018; Canales et al. 2020; Robe et al. 2020; Nikiforova et al. 2003; Wang et al. 2022). Moreover, various S-containing compounds affect plant hormone biosynthesis directly or indirectly (Wawrzynska and Sirko 2024).

Several classes of transcription factor (TF) have been identified that may co-ordinate responses to S stress. The ETHYLENE INSENSITIVE LIKE (EIL) TF family member SULFUR LIMITATION 1 (SLIM1/EIL3) is a key regulator of metabolism under S deficiency in *Arabidopsis* and tomato roots (Maruyama-Nakashita et al. 2006; Canales et al. 2020) partnering with EIL1 (Dietzen et al. 2020). Bioinformatic analysis suggested NUCLEAR FACTOR Y-A2 (NFY-A2) and the circadian regulator REVEILLE2 (RVE2) as ‘hub’ genes co-ordinating expression of S-response genes (Henriquez-Valencia et al. 2018). S availability affects expression of *Arabidopsis* R2R3-MYB family TFs that regulate biosynthesis of glucosinolates, S-containing secondary metabolites with defence functions, likely via SLIM1 (Frerigmann and Gigolashvili 2014a, b; Li et al. 2013). Moreover, S stress increases expression of a suite of *Arabidopsis* root TFs including several R2R3-MYBs; in particular, *AtMYB93* shows a sustained increase upon S deprivation and S resupply (Bielecka et al. 2014).

*AtMYB93* is part of a flowering plant-specific clade (the S24 clade) of plant R2R3-MYB transcription factors (Du et al. 2015; Gibbs et al. 2014). In *Arabidopsis*, the S24 clade consists of *AtMYB93*, *AtMYB92* and *AtMYB53* (Du et al. 2015; Gibbs et al. 2014). All three *Arabidopsis* S24 *R2R3-MYB*s show root-enriched expression with *AtMYB93* being expressed exclusively in the root (Gibbs et al. 2014). The function of the three *Arabidopsis* S24 *MYB* genes is not fully redundant. *AtMYB93*, but not *AtMYB92*, is a negative regulator of LR development and only *AtMYB93* expression is induced by auxin (Gibbs et al. 2014). As mentioned above, *AtMYB93* is significantly upregulated upon S deprivation with persistent elevated expression upon S resupply in 7-day old seedlings (Bielecka et al. 2014). *AtMYB53* is upregulated by S deprivation to a lesser extent than *AtMYB93* and is not persistently upregulated upon S resupply (Bielecka et al. 2014). *AtMYB92* shows a limited S deprivation response: upregulation in seedlings when seeds are germinated directly on S-deficient medium but not when S-replete seedlings are transferred to S-deficient medium after 8 days of growth (Nikiforova et al. 2003). Interestingly, S deprivation inhibits LR development in young *Arabidopsis* seedlings (Dan et al. 2007; Dong et al. 2019; Joshi et al. 2019) although the converse occurs in older plants (Kutz et al. 2002).

*AtMYB93* is expressed only in the root endodermis (Gibbs et al. 2014), a single-celled layer separating the outer epidermis and cortex from the inner pericycle and vasculature. The endodermis forms a selective barrier to passage of solutes, nutrients and water between the soil and the root vasculature (and hence the shoot) (Andersen et al. 2015; Geldner 2013). *AtMYB93* promoter activity is restricted to a very small number of endodermal cells overlying lateral root primordia (LRP) during early lateral root (LR) development (Gibbs et al. 2014; Shukla et al. 2021). S24 *MYB* genes and the most closely related *Arabidopsis* genes in clades S10, S11 and S42 (Du et al. 2015) are involved in synthesis of the lipophilic biopolymer suberin that surrounds mature root endodermal cells and is present in fruit and in seeds (Andersen et al. 2015; Vishwanath et al. 2015; Barberon et al. 2016). *MYB* genes act, potentially in a hierarchical fashion (Xu et al. 2022; Xu et al. 2023), to enhance suberin production. Analysis in tomato, apple and *Arabidopsis* identified a conserved cross-species gene expression ‘signature’ for suberin biosynthesis (Gou et al. 2017; Lashbrooke et al. 2016). Apple *MdMYB93* promotes biosynthesis and cell export of suberin in the fruit (Legay et al. 2016). *AtMYB39* (*SUBERMAN*), *AtMYB41*, *AtMYB53*, *AtMYB92* and *AtMYB93* are all expressed in the root endodermis and can promote suberin production (Cohen et al. 2020; Hu 2018; Kosma et al. 2014; Shukla et al. 2021; To et al. 2020; Wang et al. 2020). Lipid-, wax-, fatty acid and suberin biosynthesis genes are differentially expressed in an *Atmyb93/Atmyb92/Atmyb53* triple mutant and an *At*MYB53-overexpressing line (Klein 2019). *At*MYB41, *At*MYB93, *At*MYB53, *At*MYB92 function redundantly to promote suberin biosynthesis in the root endodermis with *At*MYB41 having the dominant role (Shukla et al. 2021). *SlMYB92* promotes suberin biosynthesis in the tomato exodermis (Canto-Pastor et al. 2024), while three *MYB*s, *OsMYB93a*, *OsMYB41* and *OsMYB76* also regulate root suberin biosynthesis in a monocot, rice (Huang et al. 2024). Expression of *AtMYB93*, *AtMYB53*, *AtMYB9* and two other MYB transcription factors is upregulated in the *kcs1-5* mutant that has impaired synthesis of very long chain fatty acids (VLCFAs) (Uemura et al. 2023), which are components of suberin (Serra and Geldner 2022).

Interestingly, an *Arabidopsis* mutant lacking all four of *At*MYB41, *At*MYB93, *At*MYB53 and *At*MYB92 has reduced suberin but has no change in lateral root development (Shukla et al. 2021), suggesting the functions of *At*MYB93 in suberin formation and root development are separable. The function of *At*MYB93 and its relatives in plant responses to S has not been investigated.

Given that *AtMYB93* has a non-redundant root function, highly specific root expression and a unique pattern of regulation by hormones and environmental signals compared to other closely related MYB genes, we hypothesise that *AtMYB93* has functions in addition to suberin biosynthesis during *Arabidopsis* root development.

## Results

### Transcriptome analysis of plants lacking *AtMYB93* reveals a potential function in responding to S

To ascertain how *AtMYB93* carries out its function, we conducted transcriptome analysis comparing gene expression in roots of wild type (Col-0) and *Atmyb93* seedlings using RNAseq

(Fig. 1; Online Resource 2). A total of 255 genes were differentially expressed (p<0.001) between Col-0 and *Atmyb93*, of which 44 were downregulated (including *AtMYB93* itself) and 211 were upregulated (Online Resource 2).

**Fig. 1.**
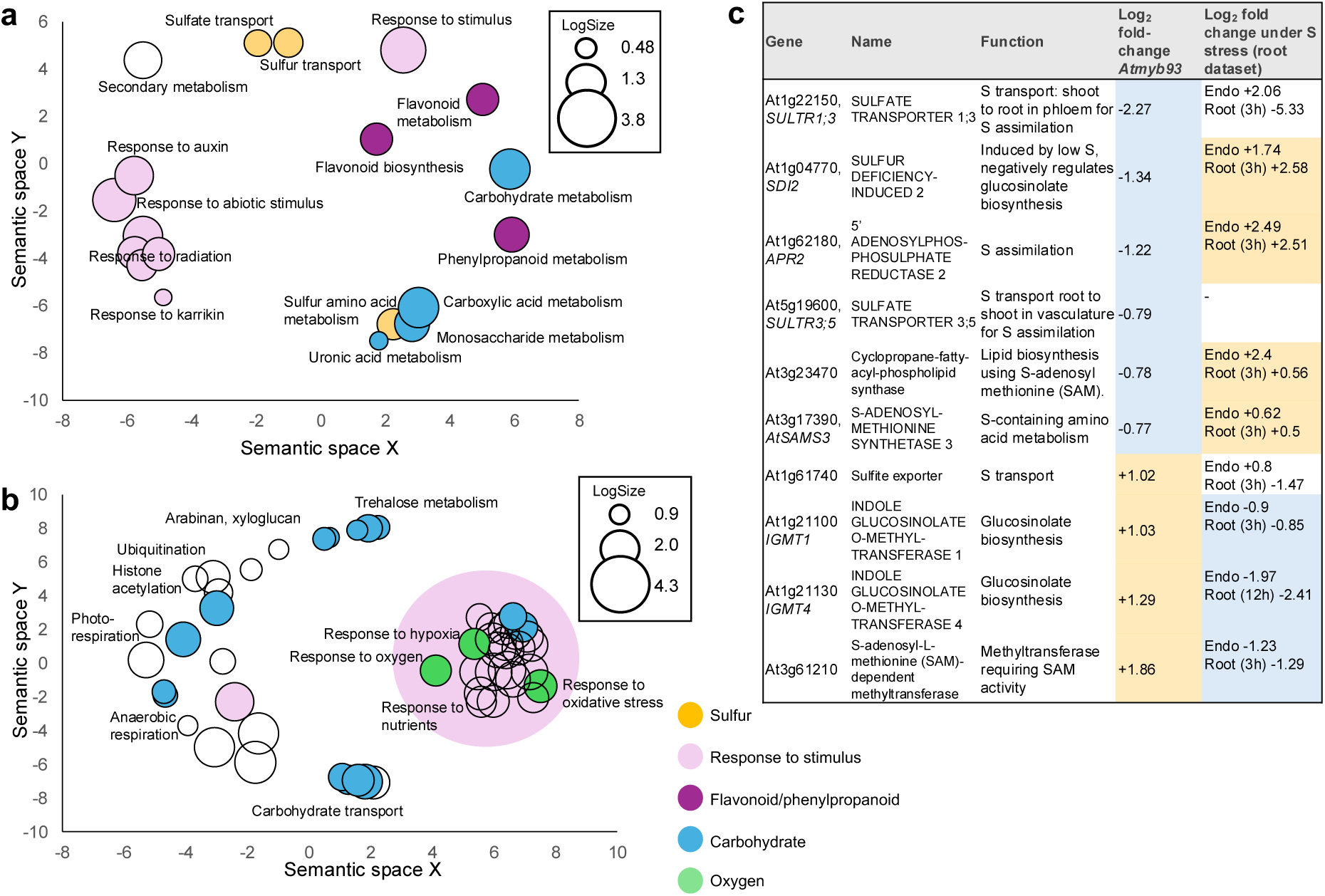
**Differentially expressed genes in the *Atmyb93* mutant are enriched for functions relating to S-, phenylpropanoid/flavonoid- and carbohydrate metabolism, and responses to a range of environmental stimuli. S-related differentially expressed genes in *Atmyb93* largely also show differential expression in response to S deprivation.** a) and b) Gene Ontology (GO) Biological Process (BP) term enrichment for genes (a) significantly downregulated (q value < 0.05) in the *Atmyb93* mutant defined by biological process and (b) significantly upregulated (q value < 0.05) in the *Atmyb93* mutant. In each panel, GO term enrichment was calculated using PlantRegMap (p<0.01) and represented with ReviGO, based on the semantic distances between GO terms. LogSize represents the log10 (number of annotations for GO Term ID in selected species in the EBI GOA database). c) Genes with S-related functions significantly differentially expressed in the *Atmyb93* mutant, showing the gene, gene name, function and log2 fold-change (down, blue and up, yellow) in *Atmyb93* or under S stress in the root. Data in the right-hand column highlights genes that also have a fold-change change >1 (down, blue or up, yellow) under S stress in either endodermis (endo, 3h S deprivation) or a whole root time series (time of largest increase shown) published by (Iyer-Pascuzzi et al. 2011) and visualized at the eFP browser (Winter et al. 2007). Two genes, At1g22150 *SULTR1;3* and At1g61740 sulfite exporter, show different regulation in the endodermis compared to the whole root

When analysed for Gene Ontology (GO) term enrichment, genes downregulated in the *Atmyb93* mutant compared to wild type showed enrichment of Biological Process (BP) terms associated with (i) S transport and metabolism, (ii) carbohydrate and sugar metabolism, (iii) phenylpropanoid and flavonoid metabolism and (iv) responses to stimuli including light, auxin and karrikin (Fig. 1a). Genes upregulated in *Atmyb93* compared to wild type were enriched for GO-BP terms associated with (i) carbohydrate (including trehalose) metabolism, transport and responses and (ii) responses to stimuli including carbohydrates and hypoxia/oxygen as well as other abiotic stimuli (Fig. 1b). These changes are reminiscent of both S deprivation responses in other studies, as outlined in the introduction to this paper, and of the transcriptome of *slim1* and *eil1* mutant plants [25].

We collated the differentially expressed genes with known functions in S transport and metabolism (Fig. 1c) and compared their differential expression in *Atmyb93* with their responses to S deprivation in whole root tissue and root endodermal tissue (Iyer-Pascuzzi et al. 2011; Winter et al. 2007). Four of the six genes with S-related functions downregulated in *Atmyb93* showed upregulation under S deprivation, while three of the four S-related genes upregulated in *Atmyb93* showed downregulation under S deprivation (Fig. 1c). Two genes, At1g22150 SULTR1;3 and At1g61740 sulfite exporter, show different regulation in the endodermis compared to the whole root (Fig. 1c).

There is no overlap between *Atmyb93* differentially expressed genes (Online Resource 2) and the largely suberin-related genes differentially expressed in the *Atmyb53/92/93* triple mutant (Klein 2019). Collectively, these data suggest that *At*MYB93, directly or indirectly, regulates genes related to S-deprivation responses in *Arabidopsis*.

### *MYB93* homologues in dicots show both root-specific expression and upregulation by S deprivation

Our previous data demonstrated that *At*MYB93 shows the most root-specific expression of *Arabidopsis* S24 clade of R2R3-MYB genes (Gibbs et al. 2014). To determine whether expression of *MYB93*-related genes outside *Arabidopsis* is root-specific, we analysed tissue-specific expression of three tomato *MYB93* homologues. The closest tomato *MYB93* homologue is *Sl04g074170* while *Sl04g056310* has previously been annotated as *SlMYB53L* (Canales et al. 2020) and *Sl05g051550* has previously been annotated as *SlMYB92* (Canales et al. 2020; Canto-Pastor et al. 2024). Note that *Sl04g074170* is different to the ‘*SlMYB93*’ identified in Kajala et al (2021), *Sl11g011050*, which is closest in protein sequence to *At*MYB48 (Kajala et al. 2021). All three *SlMYB93* homologues show root enrichment (Fig. 2a). We made transgenic *Arabidopsis* plants heterologously overexpressing *SlMYB93* (Fig. 2c) and showed that *SlMYB93* can reduce lateral root density and root length (Fig. 2c,d) similarly to *AtMYB93* overexpression (Gibbs et al. 2014).

**Fig. 2.**
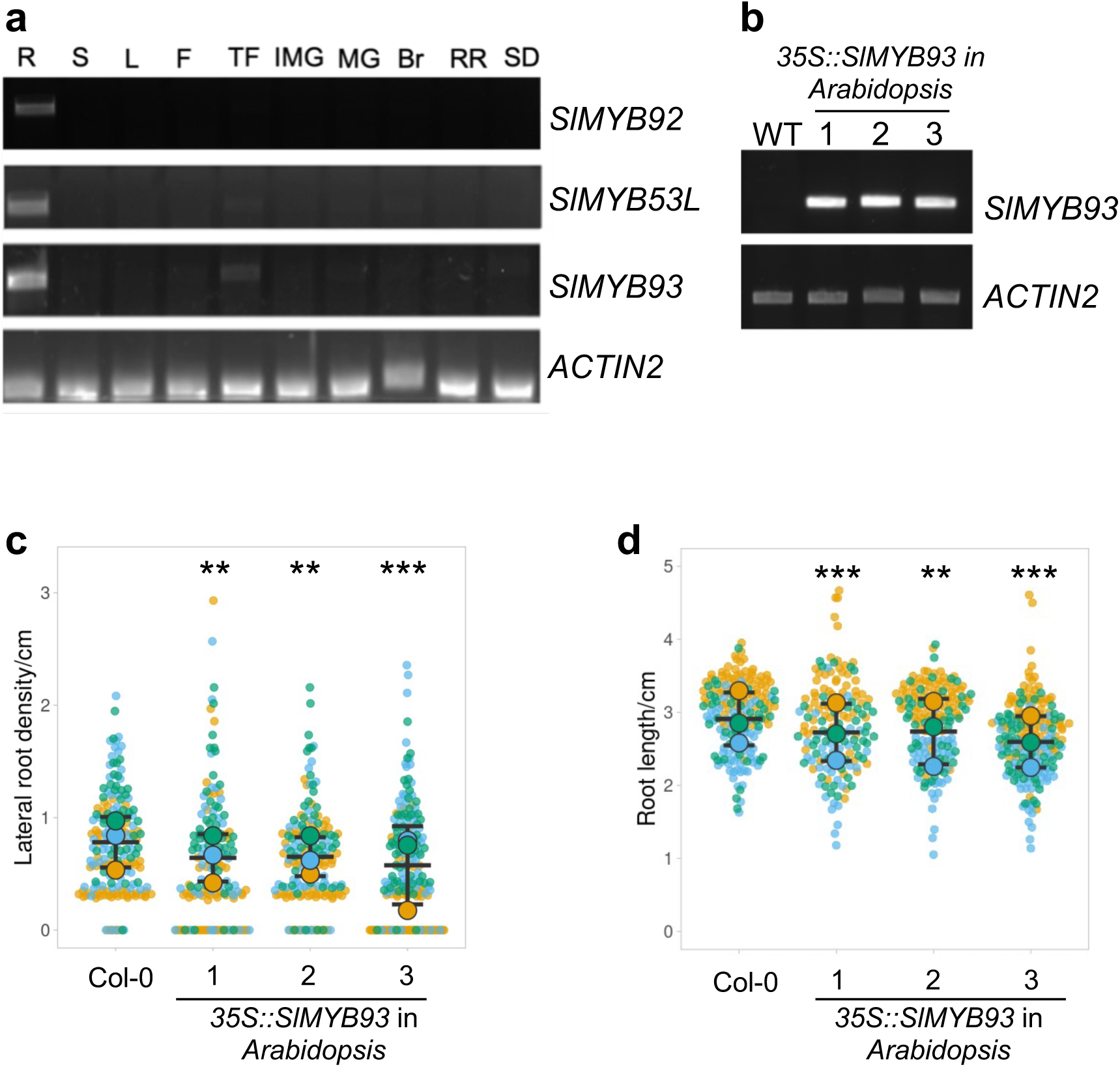
**Tomato S24-related *MYB93* homologues are root-enriched and *Sl*MYB93 functions in *Arabidopsis* similarly to *At*MYB93.** a) Root-enriched expression of tomato *MYB93* homologues demonstrated by RT-PCR. R, root; S, Stem; L, leaf; F, flower; TF, 4 days post anthesis (DPA) tomato fruit; IMG, immature green; MG, mature green; Br, breaker; RR, red ripen; SD, seed. *SlMYB93*, Solyc04g074170; *SlMYB53L*, Solyc04g056310; *SlMYB92* Solyc05g051550. The *ACTIN2* gene is shown as a control. b) RT-PCR showing heterologous expression of *SlMYB93* in *Arabidopsis* using a *p35S::SlMYB93* construct (3 independent transgenic lines) compared to a wild type control (WT, Col-0). c) Lateral root density of 3 independent transgenic *Arabidopsis* lines overexpressing *SlMYB93* (*Sl*MYB93-OX). d) Primary root length of 3 independent transgenic *Arabidopsis* lines overexpressing *SlMYB93*. 8-day old seedlings from 3 combined biological repeats are shown with data points (small coloured points) for each repeat coloured differently. Larger coloured circles represent the means of the biological repeats and black bars represent the overall mean and standard deviation of the means. In c) and d) 8-day old seedlings from 3 combined biological repeats are shown with data points (small coloured points) for each repeat coloured differently. Larger coloured circles represent the means of the biological repeats and black bars represent the overall mean and standard deviation of the means. The number of seedlings per treatment ranges from 35-75; these are the same seedlings in both panels

To extend our analysis beyond *Arabidopsis* and tomato, we analysed publicly available transcriptome data and demonstrated that root-specific or root-enriched expression of putative *MYB93* homologues is present in a wide range of, but not all, dicot species (Fig. 3). In poplar and cassava, which each have 2 closely related *MYB93* homologues, only one homologue shows root enrichment or specificity, whereas both soybean homologues show root specificity (Fig. 3). Dicot *MYB92/53*-like genes also show root enrichment (Fig. 3). Some dicot *AtMYB93*-like genes from outside the *MYB93* clade show root enrichment (*AtMYB48* and *Sl11g011050*) (Fig. 3). Some monocot *AtMYB93*-related genes show root enrichment to a lesser extent: all rice homologues (*OsMYB93a* (Os08g37970), *OsMYB93b* (Os06g17780) and *OsMYB41*(Os02g51799) (Huang et al. 2024)) are root-enriched as is one barley homologue and one maize homologue but not the genes in *Brachypodium* (Fig. 3). Overall, root specificity/enrichment is most pronounced in dicot *AtMYB93* homologues (compared to *MYB92* and *MYB53*).

**Fig. 3.**
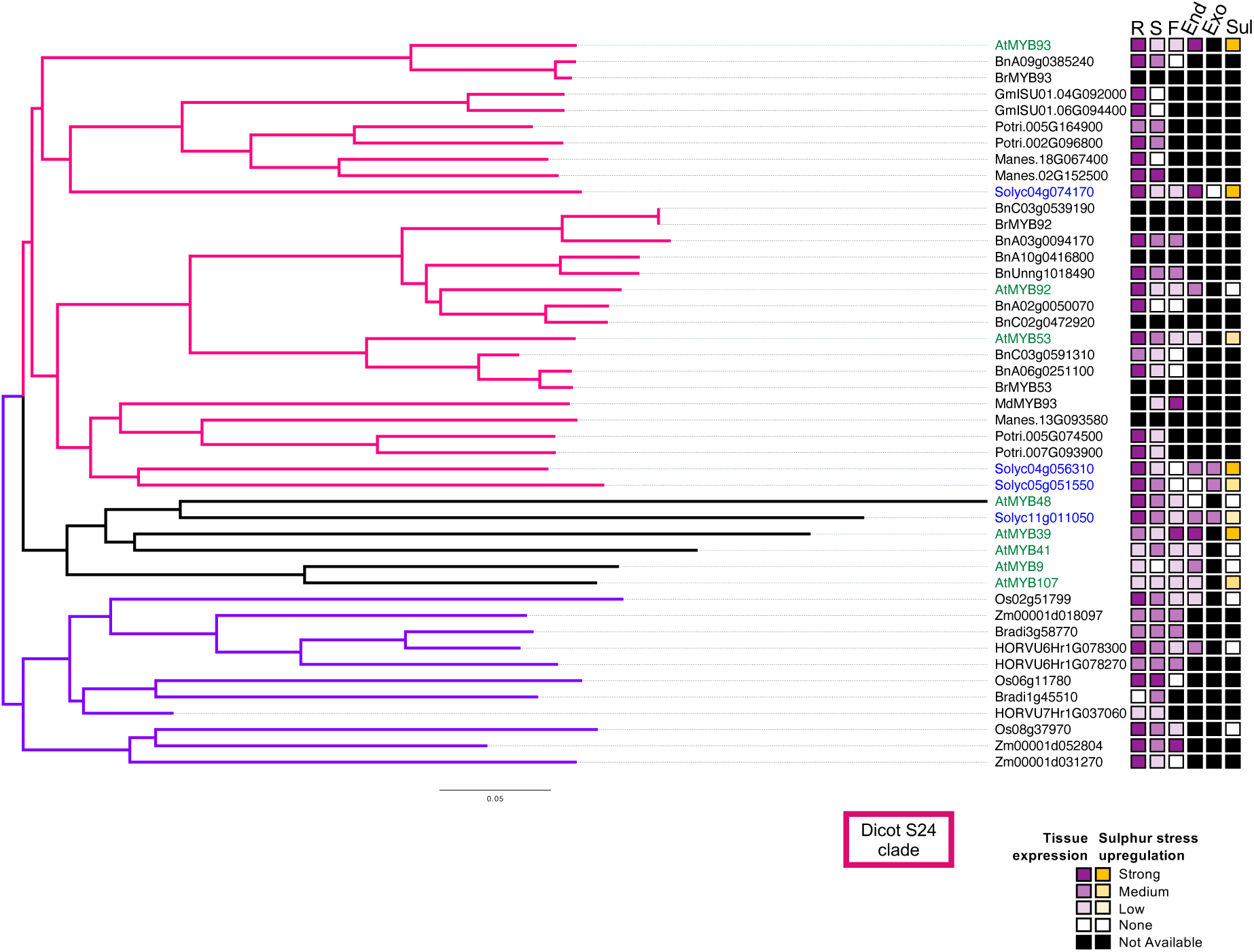
**Expression of flowering plant *MYB93* homologues: root-enrichment and dicot-specific response to S deprivation.** Phylogeny of *AtMYB93*-related genes: the clade in which the *Arabidopsis* S24 genes fall is coloured in pink, a moncot *AtMYB93*-related clade is purple and a clade containing the additional suberin-producing *Arabidopsis* MYBs is in black. *Arabidopsis* genes are shown in green and tomato genes are shown in blue. Semi-quantitative expression analysis from publicly available/published data is shown: R, root; S, shoot, F, fruit, silique or seed; End, endodermis; Exo, exodermis; Sul, S deprivation response. Pink, tissue expression; yellow, upregulation in response to S deprivation. Key to species: At, *Arabidopsis thaliana*; Bn, *Brassica napus;* Br, *Brassica rapa*; Gm, *Glycine max* (soybean); Potri, *Populus trichocarpa* (poplar); Manes, *Manihot esculenta* (cassava); Solyc *Solanum lycopersicum* (tomato); Md, *Malus domestica* (apple); Os, *Oryza sativa* (rice); Zm, *Zea mays* (maize); Bradi, *Brachypodium distachyon;* HORVU, *Hordeum vulgare* (barley). According to the notation of Huang et al (2024), Os2g51799 corresponds to *OsMYB41*, Os08g37970 corresponds to *OsMYB93a* and Os06g17780 corresponds to *OsMYB93b*

We also analysed, where available, the cell type specificity of *MYB93* homologues within the root and their response to S deprivation. Both *AtMYB93* and *Sl04g074170* show endodermal-specific and meristematic expression (Fig. 3; (Gibbs et al. 2014; Kajala et al. 2021; Waese et al. 2017)). In contrast, *Sl11g011050* is expressed in both exodermis and endodermis (Fig. 3; (Gibbs et al. 2014; Kajala et al. 2021; Waese et al. 2017)). *Sl04g056310* (*SlMYB53L*) is present in endodermis, exodermis and xylem (Fig. 3; (Gibbs et al. 2014; Kajala et al. 2021; Waese et al. 2017)) and *Sl05g051550* (*SlMYB92*) is expressed in the exodermis (Fig. 3; (Canto-Pastor et al. 2024; Kajala et al. 2021; Waese et al. 2017)). All three closest tomato *MYB93* homologues are upregulated by S deprivation (Fig. 3; (Canales et al. 2020)) with *SlMYB93* and *SlMYB53L* showing the strongest response, similarly to *Arabidopsis* (Fig. 3; (Bielecka et al. 2014)). Outside the MYB93 clade, *AtMYB107* and *Sl11g011050* also show some upregulation by S deprivation (Fig. 3; (Canales et al. 2020)). However, no change in rice *MYB93*-like genes is seen upon S deprivation (Fig. 3; (Wang et al. 2022)). Taken together, these data suggest a conserved role for dicot *MYB93*-like genes in regulating root responses to S deprivation, with the closest *MYB93* homologues specifically functioning in the endodermis.

### Changes in *MYB93* level alter plant shoot element composition

To ask whether *MYB93*’s potential link to root S-deprivation responses is associated with changes in the shoot, we first measured the elemental composition of key macronutrients (Mg, P, S, K, Ca) and micronutrients (Mn, Fe, Zn, B, Cu, Mo) in *Arabidopsis* wild type and *Atmyb93* mutant shoot tissue. Under normal growth conditions, the *Atmyb93* mutant showed generally elevated macronutrient and micronutrient levels but significant differences in S, Mg and B (Fig. 4a; Online Resource 2). Next, we investigated elemental composition of transgenic tomato lines with increased *SlMYB93* expression, *SlMYB93-OX* (Fig. 4b,c). The trends observed were in accordance with the *Arabidopsis* result, with two out of three independent *SlMYB93-OX* lines showing reduced shoot elements (Fig. 4C; Online Resource 2).

**Fig. 4.**
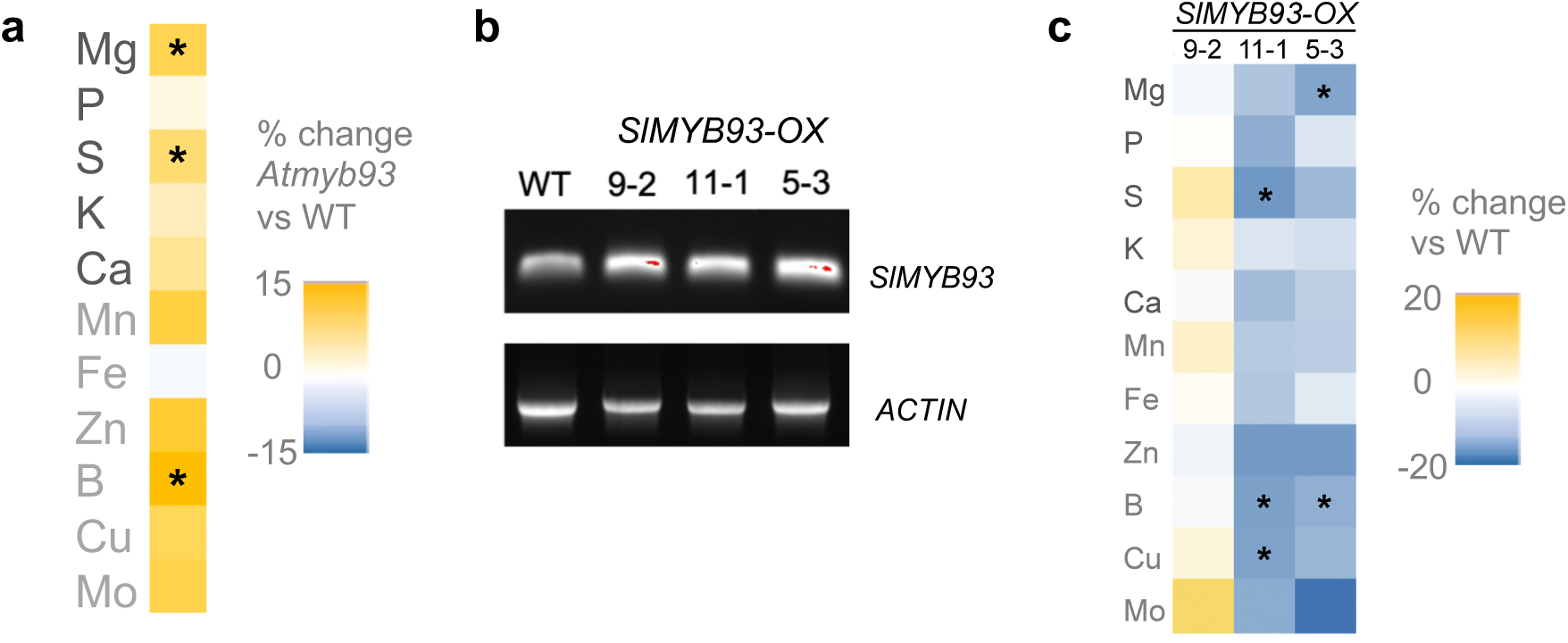
***Arabidopsis Atmyb93* mutants and tomato overexpressing *SlMYB93* show altered shoot element composition.** a) Comparison of shoot element composition in Col-0 and *Atmyb93* 21-day old plants. Heatmap shows the percentage change of means from 3 combined biological repeats. Asterisks represent significant differences (p=0.04 for each) from Mann Whitney tests on the raw data. b) RT-PCR detection of *SlMYB93* in the roots of wild type tomato plants and two independent *SlMYB93*-overexpressing lines. c) Comparison of shoot element composition in wild type and *SlMYB93-* overexpressing tomato plants. Heatmap shows the percentage change of means from 3 combined biological repeats. Asterisks represent significant differences (all p=0.04) from Mann Whitney tests on the raw data

One line showed a significant reduction in Mg and B and another showed a significant reduction in S, B and Cu (Fig. 4C; Online Resource 2). These data suggest that altering *MYB93* function in the root can lead to elemental changes particularly to S, Mg and B in the shoot.

### *Atmyb93* mutants respond to S deprivation

To further investigate the link between *At*MYB93 and plant responses to S, we compared root development responses of wild type and *Atmyb93* mutant plants under S deprivation.

A reduction in lateral root number under S starvation has previously been reported (Dong et al. 2019). A reduction in lateral root density and an increase in root length was seen in seedlings transferred to -S for 3 days after 5 days’ growth on S-replete medium that was slightly, but not significantly, less pronounced in the *Atmyb93* mutant (Fig. 5a,b). We also observed that S deprivation increases the number of adventitious roots (AR) produced at the root-shoot junction (collet) in both wild type and *Atmyb93* plants transferred to medium lacking S for 3 days after 5 days’ growth on S-replete medium, with no significant differences in AR number between wild type and *Atmyb93* being seen (Fig. 5c). We also tested the sensitivity of *Atmyb93* to O-acetyl serine (OAS), a compound that is part of the S assimilation pathway and increases in response to S deprivation (Hirai et al. 2003; Hubberten et al. 2012a; Hubberten et al. 2012b). We did not observe significant effects of OAS on either the primary root length or lateral root density of wild type or *Atmyb93* mutant plants (Fig. 5d,e).

**Fig. 5.**
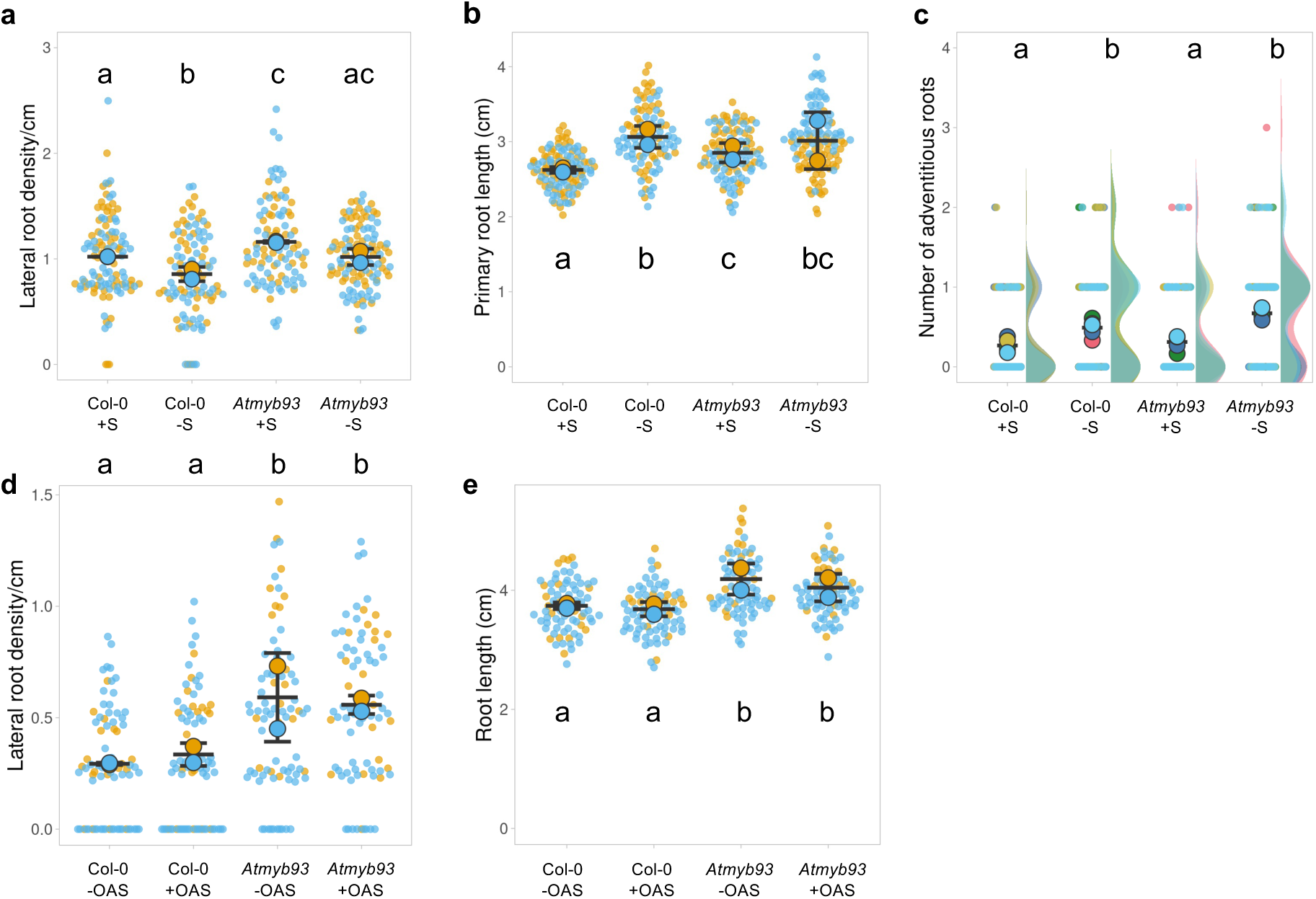
S deprivation affects root development but *Atmyb93* mutants do not show strong root or shoot phenotypes upon S deprivation or OAS treatment. a), b) Lateral root density (a) and primary root length (b) in wild type (Col-0) and *Atmyb93* seedlings grown with S for 3 days then transferred to –S medium (or +S medium control) for 5 days. 2 combined biological repeats are shown with data points (small coloured points) for each repeat coloured differently. Larger coloured circles represent the means of the biological repeats and black bars represent the overall mean and standard deviation of the means. c) Formation of adventitious roots at the collet in wild type (Col-0) and *Atmyb93* 8-day old seedlings grown for 3 days with S before transfer to –S medium (or a +S medium control). 5 combined biological repeats are shown with data points (small coloured points) and distributions (shaded areas) for each repeat coloured differently. Larger coloured circles represent the means of the biological repeats and black bars represent the standard deviation of the means. d), e) Primary root length (d) and lateral root density (e) of wild type (Col-0) and *Atmyb93* seedlings grown for 8 days on 0.1mM O-acetyl serine (OAS). 2 combined biological repeats are shown with data points (small coloured points) for each repeat coloured differently. Larger coloured circles represent the means of the biological repeats and black bars represent the overall mean and standard deviation of the means. In all panels, letters indicate differences p<0.05 in post-hoc Dunn’s test carried out after a Kruskal-Wallis test. For adventitious roots, seedling numbers range from 9 to 20 per treatment in a biological repeat. For primary/lateral root experiments ±S, seedling numbers range from 30 to 98 per treatment depending on the biological repeat. For OAS experiments, seedling numbers range from 18 to 20 per treatment in one biological repeat and 56 to 59 in the other

Collectively, these results show that S deprivation can increase adventitious root formation at the root-hypocotyl junction and suggest that, while *Atmyb93* may show slight insensitivity to S deprivation, factors in addition to *AtMYB93* also contribute to root S deprivation responses.

### S stress does not affect the spatial localisation of *AtMYB93* promoter activity

As S deprivation increases *AtMYB93* gene expression (Bielecka et al. 2014) and as *AtMYB93* transcriptional activity is restricted to a very few cells in the root, we investigated whether endodermal localization of *AtMYB93* changes under S deprivation using transgenic *Arabidopsis* expressing *pAtMYB93::GUS* (Gibbs et al. 2014). No change in the localisation and extent of *AtMYB93* promoter activity was seen under S stress (Fig. 6).

**Fig. 6.**
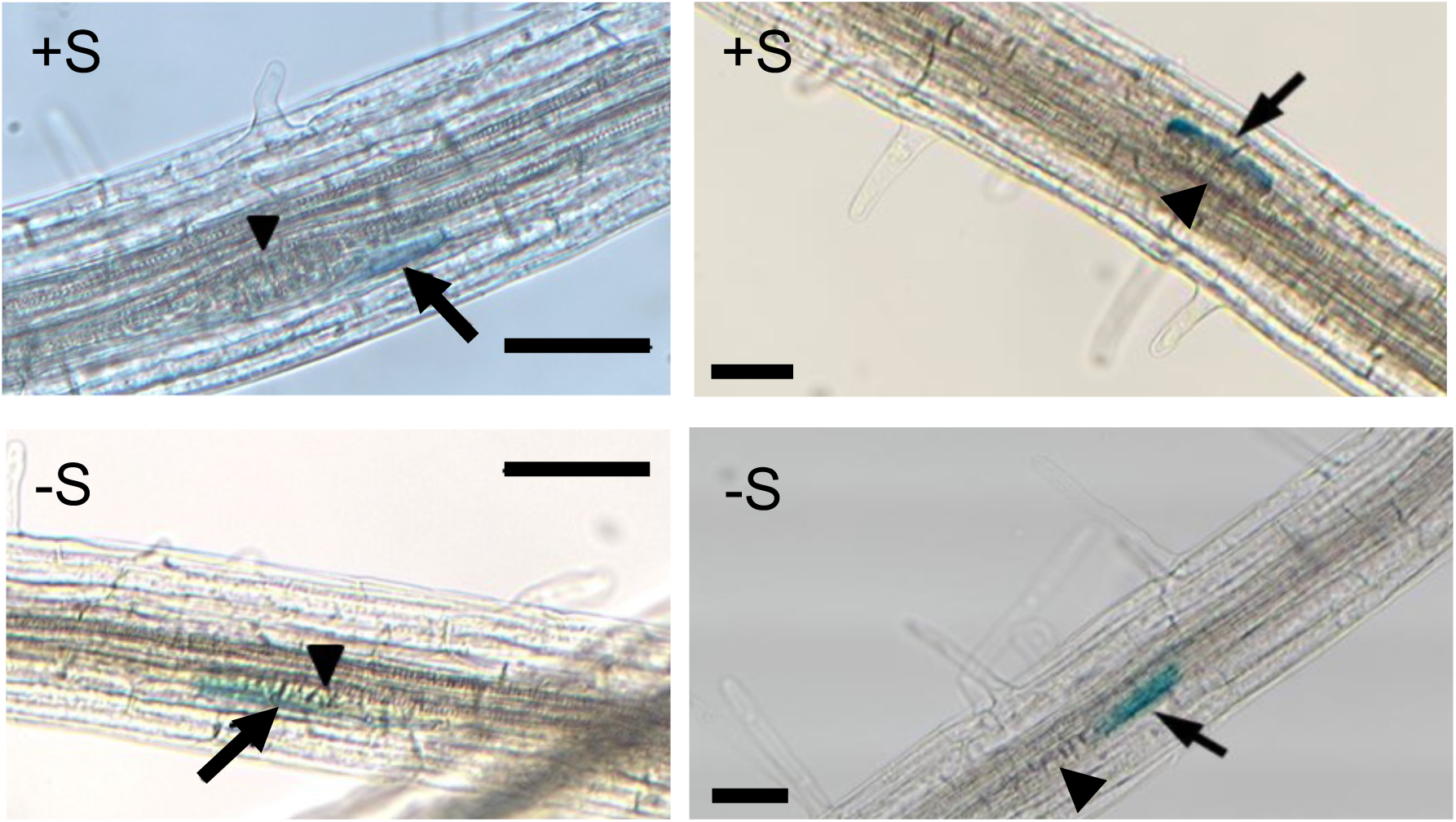
**Spatial localization of *AtMYB93* promoter activity is not changed by S stress.** *At*MYB93 promoter activity visualised in wild type plants expressing *pAtMYB93::GUS* (Gibbs et al 2014) under normal sulphur levels (+S) and S deprivation (-S) in 7-day old roots. 2 examples of each treatment are shown. Cells expressing *pAtMYB93::GUS* are highlighted with arrows. Developing Lateral Root Primordia are highlighted with arrowheads. Scale bars represent 50μm

## Discussion

### The *Atmyb93* mutant transcriptome highlights root responses to S

Despite the known (albeit redundant) role for *At*MYB93 in endodermal suberin production (Klein 2019; Shukla et al. 2021) the *Atmyb93* mutant root transcriptome does not show changes in suberin metabolism or transport genes. This is in agreement with an experiment that identified genes differentially expressed between wild type and *Atmyb93* root segments containing gravity-induced 46h lateral root primordia, which identified cell wall metabolism as an enriched biological process but no suberin metabolism genes (Uemura et al. 2023). However, there is almost no gene overlap between the dataset from root segments at this very specific, synchronised developmental time point (Uemura et al. 2023) and our whole-root dataset (Online Resource 2).

Instead, genes differentially expressed in *Atmyb93* mutant whole roots compared to wild type are enriched in GO terms for S transport and metabolism, carbohydrate metabolism, flavonoid-and phenylpropanoid metabolism and responses to oxygen and other stimuli. This profile is similar to S deprivation response transcriptomes (e.g. (Bielecka et al. 2014; Canales et al. 2020; Chandra and Pandey 2014; Forieri et al. 2017; Henriquez-Valencia et al. 2018; Lunde et al. 2008; Robe et al. 2020; Wawrzynska et al. 2022)). The trend of down- and upregulated genes suggests that the *Atmyb93* mutant is in a ‘S-replete’ state as the key S deprivation response genes *APR2* and *SDI2* are downregulated in *Atmyb93* but upregulated upon S stress (Fig. 1c; (Maruyama-Nakashita et al. 2003; Aarabi et al. 2016)). The two sulphate transporters downregulated in the *Atmyb93* mutant are the vascular transporters *SULTR1;3* (phloem) and *SULTR3;5* (xylem, pericycle and LRP), which mediate long-distance transport from source to sink and from root to shoot, respectively (Kataoka et al. 2004; Yoshimoto et al. 2003). The genomic region of At1g61740 sulfite exporter, which is upregulated in *Atmyb93*, was also identified as binding to *At*MYB93 in a DNA-affinity purification-sequencing (DAP-seq) experiment (O’Malley et al. 2016). Interestingly, root to shoot movement of sulphate under drought stress affects leaf ABA biosynthesis and stomatal closure via S-incorporation into the amino acid cysteine (Batool et al. 2018; Cao et al. 2014). As *AtMYB93* is expressed only in a very few cells of the root (Gibbs et al. 2014) it may be mediating localised changes in or responses to S levels in the root that affect responses to the environment.

### *AtMYB93*, S stress and root development

*At*MYB93 is a known negative regulator of lateral root development (Gibbs et al. 2014) that is upregulated by S stress ((Bielecka et al. 2014); this work). Previous research has shown that S stress can reduce LR development (Dan et al. 2007; Dong et al. 2019; Joshi et al. 2019) and enhance primary root elongation (Zhao et al. 2014). Although we hypothesised that these changes in root development upon S stress would be mediated by *Atmyb93*, we did not see clear differences in S deprivation root phenotypes in wild type or *Atmyb93* plants in our experiments.

Previous experiments showing LR changes in wild type plants either measured lateral root number (rather than density) and lateral root length (Joshi et al. 2019), or changes were measured only in the context of S resupply (Dong et al. 2019), or in much older (15 day) seedings (Dan et al. 2007). This suggests that S regulation of LR development may have context-dependent nuances, only some of which involve *At*MYB93, or involve *At*MYB93 working partly in concert with other related R2R3 MYBs. Moreover, S deprivation does not extend the localisation of *pMYB93::GUS* expression beyond those endodermal cells overlying lateral root primordia, similarly to what is seen with ABA (Gibbs et al. 2014), suggesting that the extent of *AtMYB93* promoter activity is tightly restricted even under stress conditions.

S stress promotes flavonoid biosynthesis and increases anthocyanin levels ((Dong et al. 2019; Bielecka et al. 2014; Nikiforova et al. 2003)). GO terms for phenylpropanoid- and flavonoid biosynthesis are enriched in the genes downregulated in the *Atmyb93* mutant compared to wild type (Fig. 1a). The key flavonoid regulator *AtMYB12* (Mehrtens et al. 2005) is downregulated In the *Atmyb93* mutant, along with genes in the flavonoid biosynthesis pathway (*CYP75B1* (a flavonoid 3’ hydroxylase), *UGT78D2*, *UGT84A1*, *UGT84A2*, *UGT90A1*; Online Resource 2). Moreover, *AtMYBL2*, an inhibitor of anthocyanin biosynthesis (Dubos et al. 2008; Matsui et al. 2008), is upregulated in the *Atmyb93* mutant (Online Resource 2). These data suggest that *Atmyb93* mutant roots would contain reduced flavonoids including anthocyanins. These gene expression changes could at least in part explain the increased root phenotypes of the *Atmyb93* mutants as flavonoids inhibit lateral root development (Brown et al. 2001; Chapman and Muday 2021) while *At*MYB12 is known to regulates cell division orientation and cell differentiation in the root (Wybouw et al. 2023).

Phenylpropanoids (which include flavonoids and lignin precursors) are required for both lignin- and suberin production and maintenance in the endodermis (Andersen et al. 2021). This could explain the role of *AtMYB93* in suberin deposition. Moreover, the transcription factor DEWAX, a negative regulator of very long chain fatty acid (VLCFA) biosynthesis (Go et al. 2014), is upregulated in the *Atmyb93* mutant (Online Resource 2) suggesting that *At*MYB93 could affect VLCFA levels surrounding the root cells in which it is expressed. However, neither the quadruple *AtMYB41/53/92/93* mutant with highly reduced suberin nor the *Atmyb92* mutant (with reduced suberin (Klein 2019)) has a lateral root phenotype (Gibbs et al. 2014; Shukla et al. 2021), arguing against a clear role for suberin itself in LR development. *AtMYB12* is downregulated in the *Atmyb39*/*suberman* mutant (Cohen et al. 2020), but the lateral root phenotype of this mutant is not known.

Interestingly, in our experiments S deprivation increases adventitious root formation at the root-hypocotyl junction. To our knowledge, this is the first time that S deprivation has been shown to affect adventitious rooting at the collet in *Arabidopsis*. S stress reduces crown root formation in rice (Grewal et al. 2018) most likely via a non-*MYB93* mechanism as *OsMYB93*s are not upregulated by S stress (Fig. 3). Flavonoids are known to inhibit AR formation in the hypocotyl in the presence of auxin (Correa Lda et al. 2012) although a mutant lacking flavonoids makes fewer ARs than wild type (Correa Lda et al. 2012). Whether the *Atmyb93* adventitious root sensitivity is linked to flavonoids is unknown.

### *AtMYB93*, shoot element composition, root architecture and root barriers

The *Atmyb93* mutant shows a general trend of elevated shoot elements, significantly S, Mg and B, while two tomato *SlMYB93*-overexpressing lines show the opposite trend. The third *SlMYB93*-overexpressing line broadly resembles wild type plants, perhaps due to position-specific insertion effects of the transgene (Schnell et al. 2015).

The role of suberin on S uptake and assimilation is not fully understood. S deprivation increases and extends suberisation in the root leading to fewer passage cells (Barberon et al. 2016; Ogden et al. 2018) and loss of endodermal suberin enhances the S deficiency phenotype of an S transporter mutant (Barberon et al. 2016) emphasising the requirement of an in-tact endodermal barrier for S uptake and retention. However, ionomic analysis shows shoot sulphate ion composition either not changing or being elevated in suberin-deficient transgenic plants made by expressing a suberin-degrading enzyme in the endodermis (*ELTP::CDEF1* and *CASP1::CDEF1* respectively, (Barberon et al. 2016)). Furthermore, another ionomic analysis shows no change in S in *ELTP::CDEF1* plants, a slight S reduction in the *Atmyb41*/*Atmyb53*/*Atmyb92*/*Atmyb93* quadruple mutant plants and no change/a slight reduction in S in plants overexpressing *AtMYB41* in the endodermis (*ELTP::MYB41* and *CDEF1::MYB41* respectively, (Shukla et al. 2021)). In both studies, the profiles of Mg and B are more in-line with our *Atmyb93* mutant data, being generally elevated with reduced suberin and reduced with elevated suberin ((Barberon et al. 2016; Shukla et al. 2021); Fig. 4). Plants with elevated suberin due to *AtMYB39* overexpression show reduced shoot elemental S (and several other elements), similarly to our *SlMYB93* overexpression data, but the *Atmyb39* mutant does not show elevated S (Cohen et al. 2020). Conversely, the enhanced suberin mutant *esb1* has elevated shoot sulphate levels (Baxter et al. 2009).

Together, these data imply that changes in suberin are not the primary driver of the changes in shoot S seen in the *Atmyb93* mutant: these changes are more readily explained by alterations in root architecture and/or changes in S assimilation and transport caused by the perturbed *Atmyb93* transcriptome. Although *SULTR3;5* is downregulated in the *Atmyb39* mutant as in *Atmyb93*, *AtSULTR3;5* and another S transporter *AtSULTR1;2* are both downregulated in an *At*MYB39-overexpressing line and there is no other S-related overlap between the *Atmyb93* and *Atmyb39* mutant transcriptomes (Cohen et al. 2020). Importantly, the fatty acid elongation required for suberin biosynthesis requires acetyl coenzyme A (acetyl coA), a S-containing compound (Woolfson et al. 2022) suggesting a possible link between suberin biosynthesis and S metabolism.

We suggest that *AtMYB93* is indirectly involved, via phenylpropanoid metabolism, in increasing the amount and extent of suberisation specifically in its highly localised regions of expression during S deprivation, whilst also additionally responding to S levels to change S transport and metabolism.

## Conclusion and future directions

We have shown that MYB93 regulates responses to S levels. The *Atmyb93* transcriptome shows a S stress signature, and MYB93 homologues in two dicot plants (*Arabidopsis* and tomato) are upregulated by S deprivation. *Arabidopsis* and tomato with perturbed *MYB93* levels show changes in shoot S content. S deprivation changes root architecture in *Arabidopsis* but this is not mediated solely by *AtMYB93*. *At*MYB93’s effect on lateral root development is likely via flavonoids rather than suberin as there is no evidence for direct suberin regulation by *At*MYB93: future work will investigate this link further.

## Materials and Methods

### Plant materials and growth conditions

*Arabidopsis* ecotype Col-0 wild type, *Atmyb93* mutant (Gibbs et al. 2014), *Atmpk3-1* mutant (SALK_151594, NASC ID N869692; (Merkouropoulos et al. 2008)) and *pAtMYB93::GUS* transgenic lines (Gibbs et al. 2014) were grown in the glasshouse in Levington M3 compost/vermiculite mix at 22°C under 16h light. For growth on plates, Col-0, *Atmyb93*, *Atmpk3* seeds were sterilised for 10 minutes in 10% Parozone^TM^ Bleach (Jeyes, Hemel Hempstead, UK) followed by 3 rinses in sterile distilled water and resuspension in 200µl distilled water. Seeds were vernalised at 4°C for 2 days in the dark and plated in rows to grow vertically at the top of half-strength Murashige and Skoog (MS) medium (M0404, Sigma-Aldrich, St Louis, Missouri, USA) pH5.6-5.8 with 1% agar.

For RNAseq analysis, the medium used was Sigma-Aldrich MS M0404 and seedlings were grown for 7 days. For root assays of Col-0, *Atmyb93* and *Atmpk3* (comparing genotypes, or OAS treatment) the medium used was half-strength MS (Sigma-Aldrich M0404), with seedlings grown for 8 days.

For S deprivation root assay experiments, the medium used was half-strength MS either containing (+S) or not containing (-S) S (MS basal salts MSP01 and MSP44; Caisson Labs, Smithfield, Utah USA) with 1% agar, with seedlings grown for 8 days. For some experiments, seedlings were initially germinated and grown on +S for 5 days and then transferred to -S for 3 days, using sterile forceps in a sterile laminar flow cabinet. All plates were sealed with micropore tape (3M, St Paul, Minnesota, USA).

For *Arabidopsis* fresh weight and shoot element analysis, Col-0 and *Atmyb93* were grown in Magenta pots (Sigma) containing 4 evenly spaced seedlings per pot. Pots were filled to half capacity (125ml) with 0.5 strength MS made using the Merck classic protocol (KGaA 2018) with 1% sucrose and 0.7% agar (Sigma-Aldrich CAS 9002-18-0). For medium without S, all sulphate ions were replaced with the chloride equivalent. Foil was wrapped round the part of the pot with agar to mimic soil darkness and facilitate root penetration of the agar. Pots were sealed with micropore tape (3M).

*Solanum lycopersicum* cultivar Micro-Tom was grown in an incubator (24°C) or glasshouse (22°C) in Levington M3 compost/vermiculite mix under 16h light. Seeds were sown directly onto compost unless transformation was to be carried out (see tomato transformation section); soil-grown plants were used for elemental analysis, RT-PCR analysis and cloning (roots were washed before making RNA).

### RNA extraction and cDNA generation

Plant tissue was ground in liquid nitrogen using RNAse-free ceramic pestles and mortars and RNA extraction was performed using an ISOLATE II Plant RNA kit (Bioline, Meridian Biosciences, Memphis, TN, USA).

### Transcriptome sequencing and analysis

RNA from ∼30mg of tissue per sample of whole 7-day old roots of wild type and *Atmyb93* (triplicate samples) was extracted and quality checked by both calculating the OD260/280 and OD 260/230 ratios of a 1µl sample using a Nanodrop analyser and by agarose gel electrophoresis to visualise ribosomal RNA integrity (Online Resource 1). Samples were sent to Novogene for further quality control (Agilent 2100 analysis), library preparation and sequencing. Briefly, sample mRNA was enriched using oligo(dT) coupled beads and RNA was fragmented using buffer containing 0.1mM ZnCl2. cDNA libraries were synthesised from mRNA fragments using random hexamers and reverse transcriptase followed by second strand cDNA synthesis using dNTPs, RNaseH and *E. coli* DNA polymerase I. The library was finished by carrying out end-repair, A-tailing, adaptor ligation, size selection and amplification by PCR. Library insert size and concentration was checked before Illumina sequencing. Raw data were transformed by base calling into sequenced reads (including sequence information and sequence quality information), which were exported in FASTQ format. For quality control, Phred scores were used to estimate the error rate of sequenced reads. Raw reads were filtered to remove adapters and low-quality reads. Cleaned FASTQ data were aligned to the *Arabidopsis* reference genome using HISAT2 (Kim et al. 2019). Clean, mapped fragments representing transcript abundance were used to estimate the expression level of each gene in each sample via fragments per kilobase of transcript sequence per million fragments paired sequenced (FKPM) values using HTseq software in union mode (Anders et al. 2015). Normalised FKPM values were subject to Pearson’s correlation analysis to estimate similarity between biological repeats. Differentially expressed genes (DEGs) were identified using DEseq2 software with the DEseq normalisation method with a false discovery rate (FDR) < 0.05. GO enrichment analysis was carried out separately on down- and up-regulated DEGs, using the GO enrichment tool in PlantRegMap (Tian et al. 2020) with a cutoff of p<0.01. Enriched GO terms were visualised using ReviGo (Supek et al. 2011).

### Phylogeny and *in silico* expression analysis of MYB93-related genes

*AtMYB93*-related protein sequences were selected as follows: (i) the *Arabidopsis* genes known to be involved in suberin formation (refs) that are (ii) also most closely

related at the protein sequence level to *At*MYB93 (Clades S24, S10, S11, S42; Du et al 2015), (iii) sequences from representative dicots and monocots identified as being most similar to *At*MYB93 by BLASTP (Altschul et al), mostly from species for which published expression data was also available. Chosen species were *Brassica napus*, *Brassica rapa*, *Glycine max* (soybean), *Populus trichocarpa* (poplar), *Manihot esculenta* (cassava), *Solanum lycopersicum* (tomato), *Malus domestica* (apple), *Oryza sativa* (rice), *Zea mays* (maize), *Brachypodium distachyon*, *Hordeum vulgare* (barley). Sequences were obtained from NCBI, Phytozome and BrassicaDB (http://brassicadb.cn/<u>#/</u>). Sequences were aligned using MUSCLE (https://www.ebi.ac.uk/jdispatcher/msa) and a simple Neighbour-Joining tree was generated, which was visualized using FigTree (http://tree.bio.ed.ac.uk/software/figtree/).

Expression data was extracted from a combination of the ePlant (Waese et al. 2017) and the *Arabidopsis* eFP browser RootII dataset (Iyer-Pascuzzi et al. 2011; Winter et al. 2007) and papers where expression of MYB93-related genes or whole-plant responses to S deprivation are recorded (Bielecka et al. 2014; Canales et al. 2020; Du et al. 2012; Jain et al. 2007; Kajala et al. 2021; Libault et al. 2009; Sekhon et al. 2011; Wang et al. 2022; Wilkins et al. 2009; Wilson et al. 2017). Data was visualised semi-quantitatively using colour intensity to provide an overall summary of trends.

### RT-PCR and semi quantitative RT-PCR

For cloning and RT-PCR in tomato, cDNA synthesis was performed from RNA using the SuperScript^TM^ III first-strand synthesis system (Invitrogen, ThermoFisher Scientific, Waltham, MA, USA). For expression analysis of *SlMYB93* homologues, PCR was carried out on the cDNA using Taq polymerase using primers in Online Resource 2.

### Generation of an *SlMYB93* overexpression construct and transformation into *Arabidopsis* and tomato

Full-length *SlMYB93* (*Solyc04g074170*) cDNA was cloned from tomato roots cv. Ailsa Craig using a proofreading DNA polymerase (Phusion^TM^, New England Biolabs, Ipswich, MA, USA) and relevant primers (Online Resource 2) into a modified pBI121 vector (Clontech), mpBI12135S, with modifications to the left and right border (Yongsheng Liu personal communication; (Cao et al. 2012)). mpBI121 contains additional restriction sites compared to pBI121, including *Kpn*I, *Sal*I and *Xho*I, and the *GUS* reporter gene has been removed, so that only the NOS terminator remains at the end of the multiple cloning site. mpBI121 contains the *nptII* kanamycin resistance gene for selection in plants. *SlMYB93* in mpBI121 was transformed into electrocompetent *Agrobacterium* strain GV3101 by electroporation at 2.5kV, 25μF capacitance and 400 Ω resistance.The *p35S::SlMYB93* construct was transformed into *Arabidopsis* via floral dip (Clough and Bent 1998).

For transformation of *p35S::SlMYB93* into tomato, *Solanum lycopersicum* cv. Micro Tom seeds were imbibed in water for 1h at 37°C and sterilised by soaking in 20% Parozone^TM^ (Jeyes, Hemel Hempstead, UK) bleach for 20 min followed by 3 washes with 30ml autoclaved water and drying on autoclaved filter paper in a laminar flow hood. Seeds were then rinsed in sterile water at least 3 further times in fresh tubes to completely remove all bleach residues and air-dried. Seeds were placed on half-strength MS basal salts (M5524, Sigma-Aldrich, St Louis, Missouri, USA) containing 2% sucrose in Magenta pots (Sigma-Aldrich, St Louis, Missouri, USA) and cold-treated (4°C) in the dark for at 2-3 days before moving to a growth room at 22°C in the dark for 2 days. If most seeds germinated, pots were transferred to a 16h light /8h dark cycle for 4-5 days. From these plants, tomato transformation was carried out as in (Cao 2022). Briefly, cotyledons were wounded and transformed with *Agrobacterium* containing *p35S::SlMYB93* before rounds of kanamycin selection to generate transformed callus producing shoots, followed by transfer to rooting medium and further transfer of rooted, transformed plants to soil.

### Shoot element analysis of *Arabidopsis* ICP-MS

*Arabidopsis* plants were grown in Magenta pots as detailed previously and harvested using tweezers at 21 days, before flowering had begun. Roots were removed with a razor blade and pooled fresh shoot tissue was heated at 60°C for 48h in an oven to remove all water. At least 100mg of dry shoot tissue for each condition was analysed by ICP-MS as in (Thomas et al. 2016). Significant differences were identified by Mann Whitney U-tests comparing *Atmyb93* mutants to wild type.

### Shoot element analysis of tomato by ICP-OES

For tomato shoot element analysis, ∼20g fresh weight of young leaves (the first 10-12 leaves behind the growing tip) were harvested from ∼6-week old plants of each genotype, which were dried at 50°C in an oven for 48h. At least 0.2g dry weight of each type of leaf tissue was sent for analysis by inductively coupled plasma optical emission spectroscopy (ICP-OES) at Yara Analytical Services, York, UK.

Briefly, samples were dried, milled, and sieved prior to analysis. Samples were digested in nitric acid using a microwave digester (CEM, Buckingham, UK). The digested samples were analysed on an Agilent ICP 5900 ICP-OES instrument (Agilent Technologies LDA UK Ltd, Stockport, UK) calibrated against standards of known concentrations. Significant differences between wild type and *SlMYB93*-overexpressing lines were identified by Mann Whitney U-tests.

### *Arabidopsis* root assay quantification

*Arabidopsis* seedlings growing vertically on +S or -S medium were photographed. Emerged lateral roots and adventitious roots (roots emerging from the collet) were counted by eye from plates and root length was measured from photographs using the freehand drawing tool in ImageJ (https://imagej.net/ij/). Lateral root density was calculated for each seedling by dividing lateral root number by primary root length.

All root data was visualised using SuperPlots ((Lord et al. 2020); https://huygens.science.uva.nl/SuperPlotsOfData/) Statistical significance was calculated via a Kruskal-Wallis test with a post-hoc Dunn’s test, or by a Mann-Whitney U-test for pairwise comparisons.

### *AtMYB93* promoter activity

Seedlings expressing *pAtMYB93::GUS* (Gibbs et al. 2014) were grown vertically on 0.5MS (KGaA 2018) plates with or without S and with 1% agar (Sigma-Aldrich CAS 9002-18-0) for 5-8 days. GUS assays were carried out as in Gibbs et al (2014).

Images were acquired by light microscopy using a GXCAM-HiChrome-S camera (GT vision, Stansfield, UK).

## Supporting information

Online Resource 1

Online Resource 2

## Funding and acknowledgements

XC was funded by China Scholarship Council studentship 201606690032. CC and HW were funded by UK Biotechnology and Biological Sciences Research Council (BBSRC) doctoral training grant BB/M01116X/1. NSG was funded by BBSRC SARIC grant, BB/N004302/1.

We thank Jasmine Carlson, Maxwell Ware, Ahmed Hussain, Lisa King, Elizabeth Chapman, Jessica Finch and Adam Elgey for related preliminary work on aspects of *At*MYB93 function and abiotic stress that informed the direction of this paper.

## Author Contributions

XC and JCC designed research. All authors performed research and analysed data. XC, HW, AO, RE and JCC visualised data. JCC supervised the project. XC and JCC wrote the manuscript. All authors reviewed the manuscript.

## Data availability

The datasets generated and analysed during this study are all included in the publication and supplemental material.

