## Supplementary material for "MYB93 regulates responses to environmental sulphur in *Arabidopsis* and tomato": Online Resource 1

**Plant Molecular Biology.**

Xulyu Cao^1,2^, Helen Wilkinson^2^, Bethany Hutton^2^, Nancy McMulkin^2^, Neil S. Graham^3^, Ross Etherington^2^, Alice Oliver^2^, Clare Clayton^2^, Harjeet Kaur^2^, Juliet C. Coates^2*^.

^1^ College of Resources and Environment, Academy of Agricultural Sciences, Key Laboratory of Efficient Utilization of Soil and Fertilizer resources, Southwest University, Chongqing, 400716, China.

^2^ School of Biosciences, University of Birmingham, Birmingham B15 2TT, UK.

^3^ Plant and Crop Sciences Division, School of Biosciences, Sutton Bonington Campus, University of Nottingham, LE12 5RD, UK.

### ^*^Corresponding author. ORCID 0000-0002-2381-0298

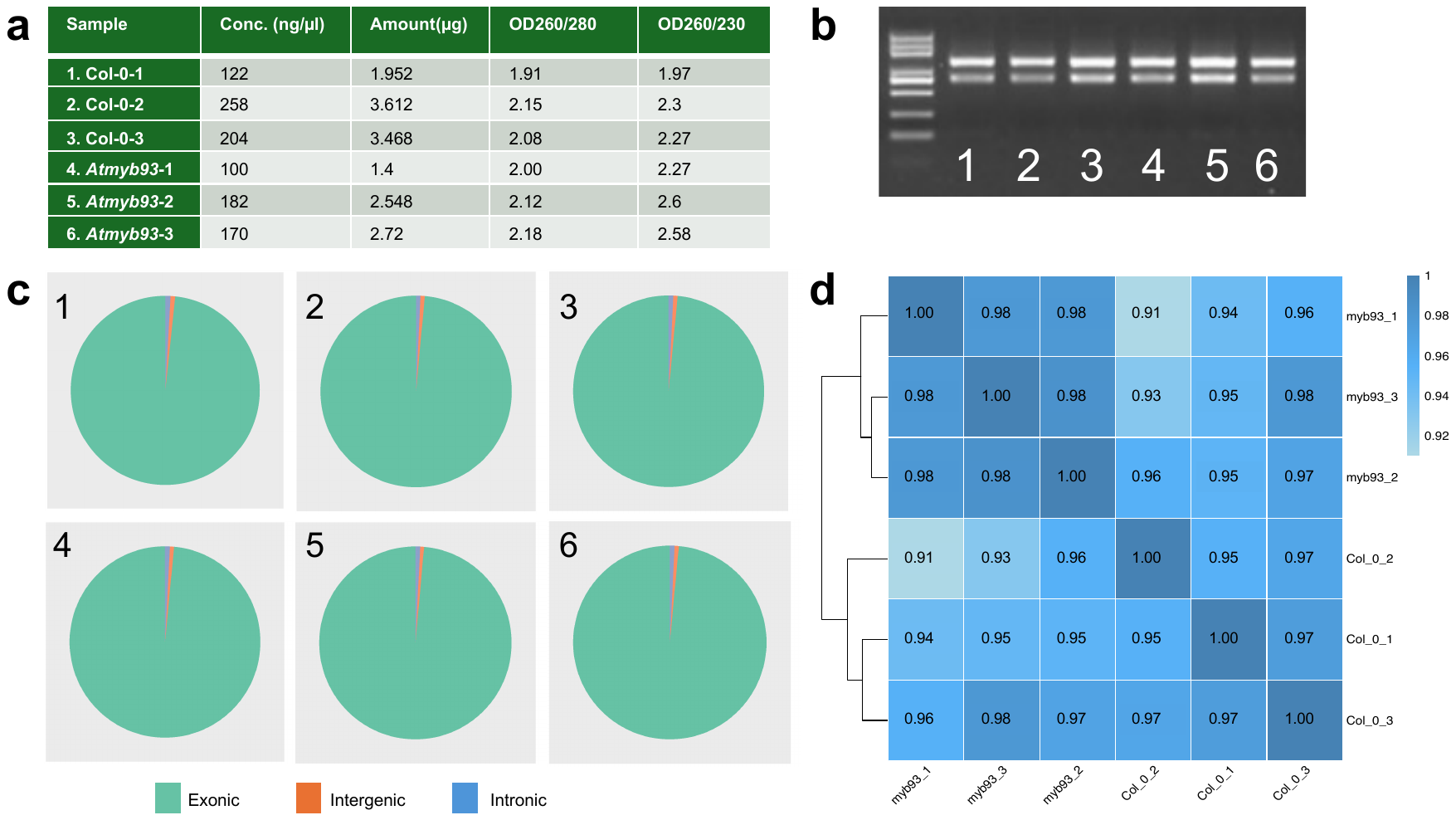
**Online Resource 1. Starting material used for RNAseq analysis and quality control of the RNAseq experiment.**

a) Concentrations and OD/OD ratios for each RNA sample: 3 wild type and 3 Atmyb93. Whole roots were harvested 7 days after germination.

b) Sample of each RNA preparation run on an agarose gel. Sample numbers are as in A), 1kb ladder is shown on the left. 28S and 18S ribosomal RNA (rRNA) bands are visible for each sample.

c) FASTQ data aligned to reference genome using HISAT2 with percentage of reads mapped to genome regions. >98% of mapped regions for each sample are classified as exons, while <2% of mapped sequences were from introns or intergenic regions. Phred scores in FASTQ format were used to estimate the error rate of sequenced reads for quality control. All samples were considered high quality with an error rate of 0.0004. Sample numbers as in A).

d) Heatmap showing Pearson’s correlation analysis on normalized FKPM data (18241 annotated genes). Correlation coefficient values of >0.9 exist between biological repeats. All Col-0 samples cluster together and all Atmyb93 samples cluster together, showing repeatability of the experiment and differences between the two genotypes.
